# Prion protein-derived cell-penetrating peptide inhibits type II diabetes-associated islet amyloid polypeptide aggregation and cytotoxicity

**DOI:** 10.1101/2025.10.09.681467

**Authors:** Yujeong Oh, Palanikumar Loganathan, Madeline Howarth, Debabrata Maity, Morad Mustafa, Sunil Kumar, Andrew D. Hamilton, Mazin Magzoub

## Abstract

Islet amyloid polypeptide (IAPP) is a 37-residue peptide hormone co-packaged and co-secreted with insulin by pancreatic β-cells. A pathological hallmark of type II diabetes is the self-assembly of IAPP into β-sheet rich amyloid fibers, which is associated with β-cell impairment. Previously, we showed that a cell-penetrating peptide (CPP) construct, consisting of a hydrophobic signal sequence coupled to a polycationic nuclear localization signal (NLS)-like sequence, exhibited potent anti-prion activity and antagonism of Alzheimer’s disease-associated amyloid-β (Aβ) peptide aggregation and neurotoxicity. Here, we have extended this approach towards type II diabetes by assessing the efficacy of the CPP construct, designated as NCAM1-PrP, in inhibiting IAPP oligomerization, fiber formation and associated cytotoxicity. Using complementary *in vitro* and *in silico* experiments, we show that NCAM1-PrP effectively modulates IAPP’s toxic structures into non-toxic conformations. This study underlines the potential of our designed CPP-based therapeutic approach as a versatile tool in the battle against amyloid-associated pathologies.

## INTRODUCTION

Many globally prevalent degenerative disorders are associated with misfolding and accumulation of specific proteins or peptides into abnormal aggregates termed amyloids.^1^ Prominent amongst these disorders are Alzheimer’s disease (AD), Parkinson’s disease (PD), prion diseases or transmissible spongiform encephalopathies, and type II diabetes (T2D), which are mediated by the self-assembly of the amyloid-β peptide (Aβ) and tau protein, α-synuclein (α-syn), prion protein (PrP), and islet amyloid polypeptide (IAPP), respectively.^1^ These proteins or peptides are usually monomeric, soluble and intrinsically disordered in their native or physiological conformation.^2^ In amyloidosis, these monomeric precursors first self-assemble into oligomers and then fibrils, which are characterized by a canonical cross-β structure, high thermodynamic stability and resistance to proteolytic digestion.^2^ While the deposition of these aggregates into extracellular plaques or intracellular inclusions is a hallmark of amyloid diseases, the consensus is that soluble oligomeric intermediates are the primary source of toxic gain of function.^3,4^

IAPP (also known as amylin) is a 37-residue, natively unstructured, peptide hormone that is co-packaged and co-secreted with insulin by pancreatic β-cells in the islets of Langerhans.^5^ IAPP plays an important role in helping to maintain glucose homeostasis, which the peptide does by a combination of slowing gastric emptying and promoting satiety.^5^ However, IAPP is highly aggregation-prone, and the peptide’s self-assembly and the associated toxic gain-of-function are strongly implicated in β-cell death in T2D.^6^ Although it has long been held that this amyloidogenic behavior is driven by the central hydrophobic core, IAPP_20–29_ (Table 1) – which is the segment that not only has the highest propensity for aggregation in isolation, but also exhibits the greatest sequence variation between species in which IAPP can and cannot form amyloid – it has become apparent that other regions strongly contribute to the peptide’s pathogenic self-assembly.^7–9^ Of note, a ∼22-residue membrane-binding N-terminal subdomain is important for increasing the nucleation potential of IAPP_20–29_.^10,11^ This catalysis is proposed to be due to interactions of the N-terminal subdomain raising the effective local concentration and relative orientation of IAPP_20– 29_.^10,12^ Thus, the aggregation of IAPP is not governed by a single molecular motif; rather, it is a more complex process, yielding polymorphic and dynamic oligomeric intermediates.^6^

**Table 1.**
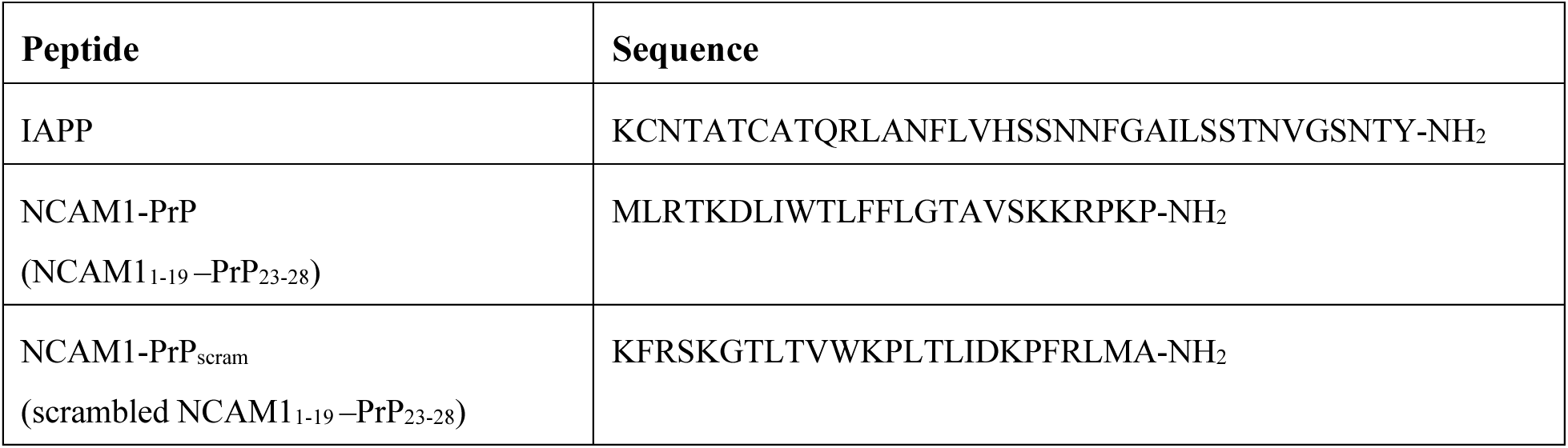
Primary sequences of IAPP and the designed amyloid inhibitor cell-penetrating peptide (CPP) construct, NCAM1-PrP, and its scrambled variant NCAM1-PrPscram.

Given its important physiological roles, interfering with the presence of IAPP will likely lead to detrimental downstream effects. Thus, a more viable therapeutic strategy for T2D is identifying or developing therapeutic agents that effectively target the peptide’s toxic oligomers. This is often achieved using synthetic small molecules, peptides, macrocyclic hosts and antibodies that either stabilize IAPP in a monomeric state thereby preventing its pathogenic self-assembly, dissociate or clear the peptide’s toxic oligomers, or remodel the oligomers into nontoxic off-pathway aggregates.^13–20^

A promising class of amyloid inhibitors are cell-penetrating peptides (CPPs) derived from PrP.^21,22^ CPPs are short peptides (∼5–40 residues) that enter cells with high efficiency and low toxicity *in vitro* and *in vivo*.^23^ The original PrP-derived CPP consists of two segments: a hydrophobic signal sequence (PrP residues 1–22) followed by a polycationic nuclear-localization signal (NLS)-like sequence (PrP residues 23–28: KKRPKP).^24,25^ NLS sequences are found in ∼90% of all nuclear proteins and usually direct their proteins to the cell nucleus.^26^ However, occasionally NLS-like sequences, which are typically hexapeptides with at least four positively charged residues, are found in non-nuclear proteins such as PrP.^27^ The N-terminal NLS-like sequence of PrP is critical for the protein’s internalization in its recycling between the cell surface and endosomal compartments.^28–30^ Yet the NLS-like sequence alone is not readily internalized into cells and requires a hydrophobic counterpart, such as the signal sequence, in order to acquire CPP properties.^31^

Treatment of prion-affected neuronal hypothalamic cells with the PrP-derived CPP effectively reduced the levels of disease-associated scrapie isoform (PrP^Sc^), without affecting endogenous PrP^C^ levels.^21^ Additionally, the CPP substantially delayed infection of healthy neuronal hypothalamic cells exposed to scrapie.^21^ These intriguing results prompted extensive structure-activity studies in order to optimize the CPP design, which yielded the following findings: (i) truncating the signal peptide (PrP_1–22_) lead to a loss of the anti-prion activity of the CPP; (ii) likewise, replacing PrP_1–22_ with conventional cationic or hydrophobic CPPs (i.e. TAT_48-60_, penetratin or transportan-10) abolished the anti-prion effect; (iii) on the other hand, replacing PrP_1-22_ with a shorter hydrophobic signal sequence from another plasma membrane-anchored glycoprotein like PrP, the neural cell adhesion molecule-1 (NCAM1_1–19_), enhanced the anti-prion potency of the CPP.^31^

The inhibition of prion conversion was attributed to NLS-like sequence of the PrP-derived CPPs, and given that this polycationic sequence has been shown to be a ‘general amyloid marker’ that selectively binds to a wide range of amyloid oligomers and fibrils via interactions with a common supramolecular feature of protein aggregates, this suggested that the PrP_NLS_-based CPPs could potentially inhibit the pathogenic self-assembly of other amyloid proteins.^31,32^ Subsequently, we demonstrated that PrP_NLS_ coupled to NCAM1_1–19_ (NCAM1-PrP; Table 1) effectively stabilized Aβ in a non-amyloid state and protected neuronal cells against Aβ-induced cytotoxicity.^22^ In the present study, we have extended our CPP-based approach towards T2D by investigating the effects of NCAM1-PrP on IAPP amyloid self-assembly and the downstream toxic effects.

## RESULTS

### Effect of NCAM1-PrP on IAPP amyloid aggregation

The capacity of the NCAM1-PrP CPP construct to modulate IAPP self-assembly was first evaluated using the thioflavin-T (ThT) based amyloid kinetic assay. ThT is a small molecule probe that exhibits a substantial increase in fluorescence intensity upon selectively binding to the cross-β structure of amyloid fibrils.^33^ Amyloid aggregation typically occurs in three stages: an initial lag phase primarily driven by nucleation of soluble misfolded protein, a rapid elongation phase where the subsequent nuclei aggregate and form ThT-positive fibrils, and a final plateau phase where soluble peptides and fibrils reach equilibrium.^34^ This nucleation-dependent process yields a characteristic sigmoidal ThT curve from which the amyloid aggregation kinetics can be quantified.^34^

IAPP alone exhibited a sigmoidal ThT curve, indicating amyloid aggregation, with a t_50_ (time required to reach 50% of maximum ThT fluorescence intensity) of 2.3 ± 0.2 h (Figure 1a). This is within the range reported for IAPP under similar conditions.^16^ NCAM1-PrP antagonized IAPP aggregation in a dose-dependent manner, with complete inhibition observed at an equimolar ratio of the CPP (Figure 1b). The ThT assay results were visually validated using transmission electron microscopy (TEM). IAPP was incubated alone or co-mixed with an equimolar ratio of NCAM1-PrP at 37 °C for 24 h. While extensive fibrils were observed in samples of IAPP alone, these were completely absent in the co-mixed samples (Figure 1b), confirming that NCAM1-PrP abolishes IAPP fibrillation. As a control, we monitored aggregation of IAPP co-mixed with a scrambled sequence of the CPP (NCAM1-PrP_scram_). In marked contrast to NCAM1-PrP, treatment with NCAM1-PrP_scram_ accelerated IAPP amyloid aggregation (t_50_ = 1.33 ± 0.2 h) (Figure 1e). This underlines the importance of the CPP construct’s primary structure – a hydrophobic signal sequence followed by a polycationic NLS-like segment – for its amyloid inhibitory capacity.

**Figure 1.**
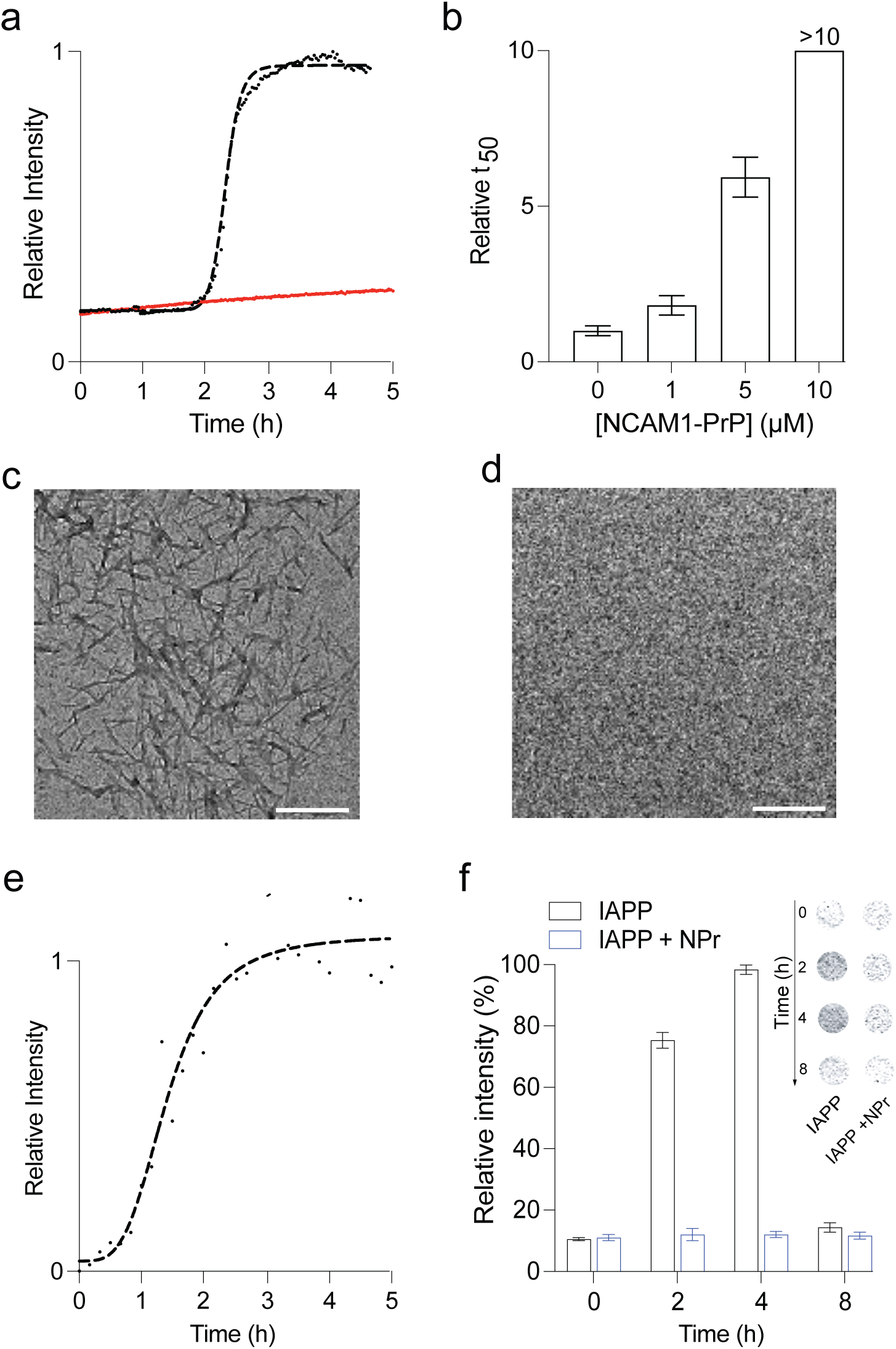
Effects of NCAM1-PrP on IAPP oligomerization and amyloid formation. (**a**, **b**) Thioflavin T (ThT)-based kinetic amyloid aggregation profiles of 10 μM IAPP in the absence (black) or presence (red) of NCAM1-PrP at an equimolar ratio (**a**), along with the relative t_50_ values of 10 μM IAPP co-mixed with increasing concentrations (0–10 μM) of NCAM1-PrP (**b**) (*n* = 3). (**c**, **d**) Transmission electron microscopy images of 10 μM IAPP incubated alone (**c**), or co-mixed with NCAM1-PrP at an equimolar ratio (**d**), for 24 h. (**e**) Kinetic aggregation profile of 10 μM IAPP in presence of the scrambled CPP sequence (NCAM1-PrP_scram_) at an equimolar ratio. (**f**) Dot blot assay to detect IAPP oligomers. Plot of the chemiluminescence signal intensity calculated from the dot blot assay (*inset*) for samples of 5 μM IAPP incubated alone or with NCAM1-PrP (NPr) at an equimolar ration for the indicated durations and subsequently treated with the oligomer-specific polyclonal antibody (A11).^3^ Error bars represent the SEM of at least 5 independent trials.

Next, we probed the effects NCAM1-PrP on IAPP oligomerization using the dot blot immunoassay. IAPP was incubated alone, or with an equimolar of NCAM1-PrP, for 0–8 h and then detected using an amyloid-oligomer specific polyclonal antibody (A11).^3^ For IAPP alone, the chemiluminescence signal intensity increased over the first 4 h, indicating formation of soluble pre-amyloid oligomers of the peptide, but subsequently decreased at 8 h, reflecting conversion of the oligomers to fibrils (Figure 1f). However, in the presence of NCAM1-PrP, the intensity did not increase above background throughout the time course of the experiment, which signifies a lack of IAPP oligomer formation (Figure 1f). Taken together, the aggregation assays show that the NCAM1-PrP CPP construct effectively inhibits IAPP oligomerization and subsequent fibril formation.

### The NCAM1-PrP CPP construct rescues IAPP-induced cytotoxicity

The effects of NCAM1-PrP on IAPP cytotoxicity were probed in RIN-m rat insulinoma cells, which have established use in studies of IAPP trafficking, fibrillation and toxicity.^16,35^ Viability of RIN-m cells exposed to IAPP, alone or in the presence of NCAM1-PrP, was assessed using the CellTiter 96 AQueous One Solution assay, which quantifies reduction of the tetrazolium compound MTS (3-(4,5-dimethylthiazol-2-yl)-5-(3-carboxymethoxyphenyl)-2-(4-sulfophenyl)-2H-tetrazolium, inner salt) to soluble formazan in living cells by mitochondrial NAD(P)H-dependent dehydrogenases.^36,37^

Cytotoxicity of IAPP scaled with peptide concentration and incubation time. Specifically, treatment with 5 or 10 μM IAPP for 24 h decreased viability of the RIN-m cells to 60±3 or 51±3% of controls using protein-free carrier, respectively, while prolonging exposure of the cells to 5 μM IAPP to 96 h reduced cell viability further to 22±4% (Figure 2a,b). Co-mixing with NCAM1-PrP rescues IAPP-induced cytotoxity in a concentration-dependent manner, with an effective concentration (EC_50_) of 1.2±0.1 μM (Figure 2c). Importantly, complete inhibition of IAPP cytotoxicity was observed at a molar ratio of 2:1 (IAPP:NCAM1-PrP) (Figure 2c). Together with the aggregation assays (Figure 1), these results suggest that NCAM1-PrP forms complexes with IAPP that prevent its amyloid self-assembly and the associated cytotoxicity.

**Figure 2.**
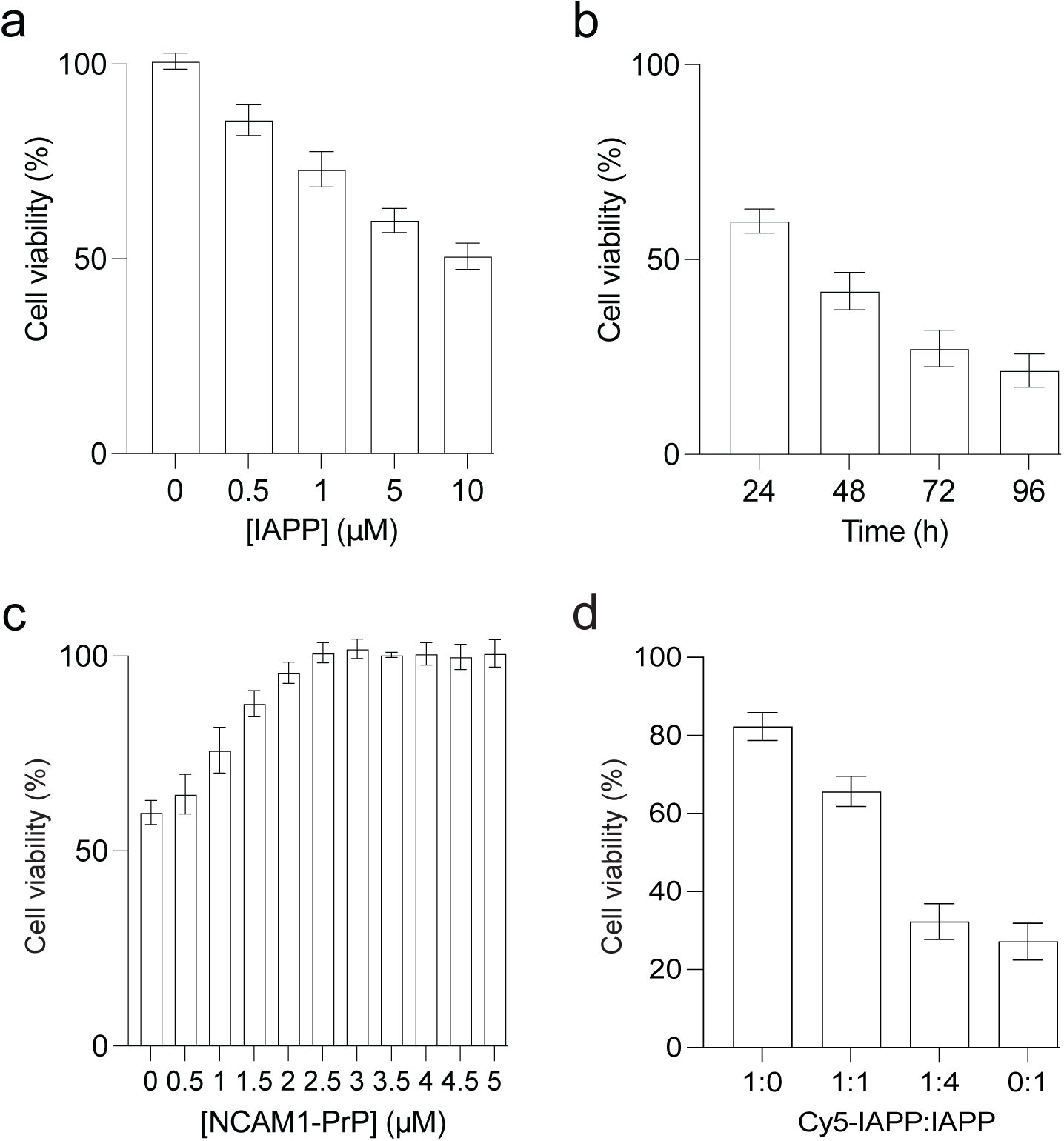
Effects of NCAM1-PrP on IAPP cytotoxicity. **(a**–**c)** Viability of RIN-m cells treated with the indicated concentrations of IAPP for 24 h (**a**), with 5 μM IAPP for the indicated durations (**b**), or with 5 μM IAPP co-mixed with increasing concentrations of NCAM1-PrP for 24 h (**c**). (**d**) Viability of RIN-m cells treated with incubated with 5 μM mixtures of IAPP and Cy5-labeled IAPP (Cy5-IAPP) at the indicated molar ratios for 72 h. Cell viability was assessed using the MTS assay, with the % viability determined form the ratio of the absorbance of the treated cells to the control cells (*n* = 3). Error bars represent the SEM of 4 independent triplet-well trials.

In order to shed the interactions of IAPP and NCAM-PrP in a cellular milieu, we monitored the cellular uptake and intracellular localization of the peptides, alone and co-mixed, using confocal fluorescence microscopy. IAPP and NCAM1-PrP were N-terminally labelled with red Cy-5 and green Alexa Fluor 488 (A_488_), respectively. Since tagging short amyloid peptides with a fluorophore can inhibit their self-assembly and associated cytotoxicity,^38–40^ here we doped unlabeled IAPP was with Cy5-labeled IAPP (Cy5-IAPP) at a 4:1 molar ratio, a mixture which exhibits the same behavior as the unlabeled peptide (Figure 2d). Treatment of RIN-m cells with IAPP/Cy5-IAPP or A_488_-NCAM1-PrP for 24 h resulted in localization of the peptides to lysosomes and, to a lesser extent, mitochondria (Figure 3a). This indicates that cellular internalization of the peptides occurs primarily by endocytosis, with subsequent escape from endocytic compartments and localization to mitochondria.

**Figure 3.**
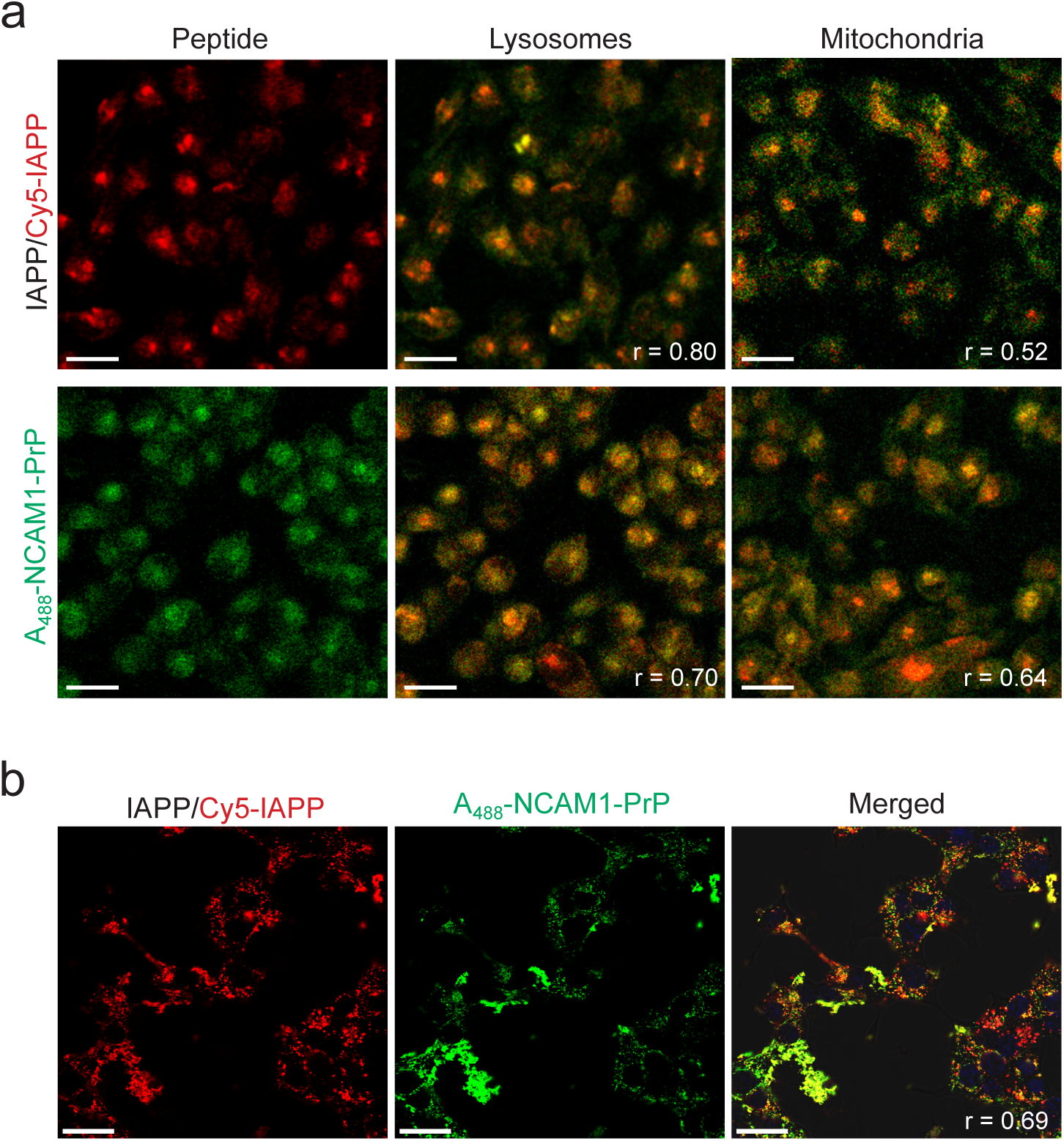
Interaction of IAPP and NCAM1-PrP in a cellular environment. (**a**) Intracellular localization of IAPP and NCAM1-PrP. RIN-m cells were treated with 5μM of IAPP/Cy5-IAPP (4:1 molar ratio) or 5μM of Alexa488-labeleled NCAM1-PrP (A_488_-NCAM1-PrP) for 24 h prior to imaging. Cells were co-stained with DAPI and LysoTracker (Red DND-99 or Green DND-26) or MitoTracker (Red FM or Green FM). (**b**) Cellular uptake of IAPP in the presence of NCAM1-PrP. Cells were incubated with 5μM of IAPP/Cy5-IAPP (4:1 molar ratio) co-mixed with A_488_-NCAM1-PrP at an equimolar ratio for 24 h prior to imaging. Colocalization was quantified using the Pearson correlation coefficient (r), which measures pixel-by-pixel covariance in the signal level of two images.^74^ Scale bar = 20 μm.

Simultaneous addition of IAPP/Cy5-IAPP and A_488_-NCAM1-PrP at an equimolar ratio to RIN-m cells yielded intracellular and extracellular complexes of the two peptides (Figure 3b). Importantly, pre-treatment of RIN-m cells with Cy5-IAPP/IAPP for 24 h, followed by addition of A_488_-NCAM1-PrP at an equimolar ratio and incubation of the cells for an additional 48 h, again resulted in strong co-localization of the two peptides intracellularly and, to a lesser extent, extracellularly (Figure 4b). Moreover, the time-delayed addition of NCAM1-PrP rescued IAPP-induced cytotoxicity: exposure of the cells to 5 μM IAPP for 72 h reduced viability to 27±5%, with a 6 h delayed addition of an equimolar amount of NCAM1-PrP fully restoring cell viability, while even prolonged delays in addition of the CPP to 12 and 24 h post IAPP treatment recovering viability to 94±4 and 85±6%, respectively (Figure 4a). Thus, the CPP property of NCAM1-PrP allows it to target both intracellular and extracellular IAPP and effectively inhibit IAPP-mediated cytotoxicity.

**Figure 4.**
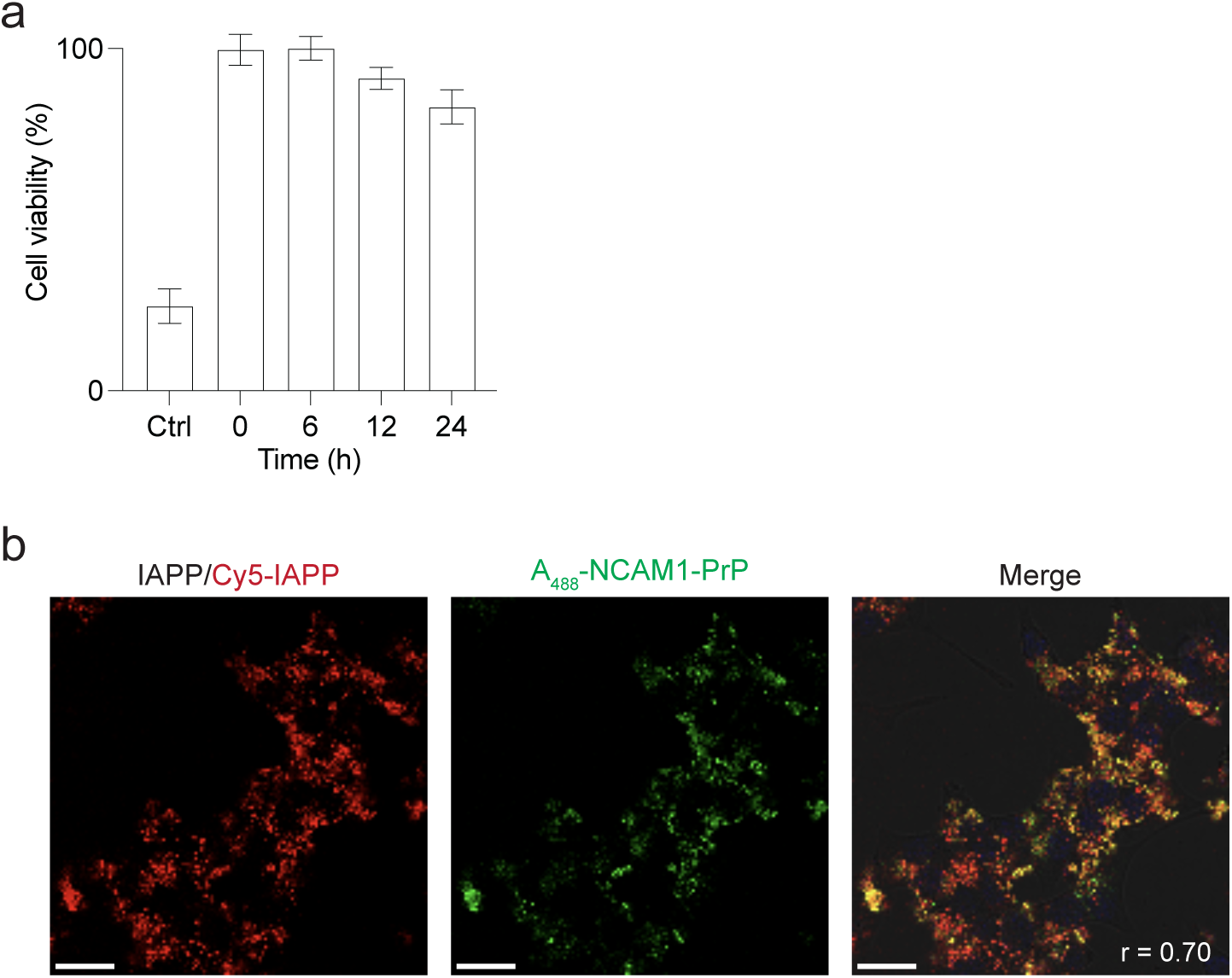
Targeting of intracellular IAPP by NCAM1-PrP. (**a**) Effects of time-delayed addition of NCAM1-PrP on IAPP cytotoxicity. RIN-m cells were pre-treated with 5 μM IAPP, followed by addition an equimolar amount of NCAM1-PrP at the indicated time points (0–24 h) post IAPP treatment. The cells were then maintained for a total incubation time with peptides of 72 h, and cell viability was quantified using the MTS assay, with the % viability determined form the ratio of the absorbance of the treated cells to the control cells (*n* = 3). Error bars represent the SEM of 4 independent triplet-well trials. (**b**) Effects of time-delayed addition of NCAM1-PrP on intracellular localization of IAPP. RIN-m cells were pre-treated with 5μM of IAPP/Cy5-IAPP (4:1) for 24 h, followed by addition of 5μM of A_488_-NCAM1-PrP and incubation of the cells for another 48 h prior to imaging. Colocalization was quantified using the Pearson correlation coefficient (r). Scale bar = 20 μm.

### Characterizing the IAPP-CPP binding interaction

In order to determine the binding site(s) of NCAM1-PrP on IAPP, we employed two-dimensional heteronuclear single quantum coherence (HSQC) NMR of recombinant IAPP (with an amide-free C-terminus) co-mixed with the CPP at an equimolar. Chemical shift assignments of ^15^N-labeled IAPP residues were based on published reports.^20,41^ We observed perturbation of residues in the N-terminal domain (Thr4–Asn22) of IAPP upon addition of NCAM1-PrP, while residues towards the C-terminus of the amyloid peptide were largely unchanged (Figure 5). Specifically, peak volumes for many of the N-terminal residues decreased noticeably, with the greatest change occurring at His18, where the peak disappeared completely (Figure 5). These results suggest that NCAM1-PrP modulates IAPP’s fibrillation properties by binding to its N-terminal subdomain.

**Figure 5.**
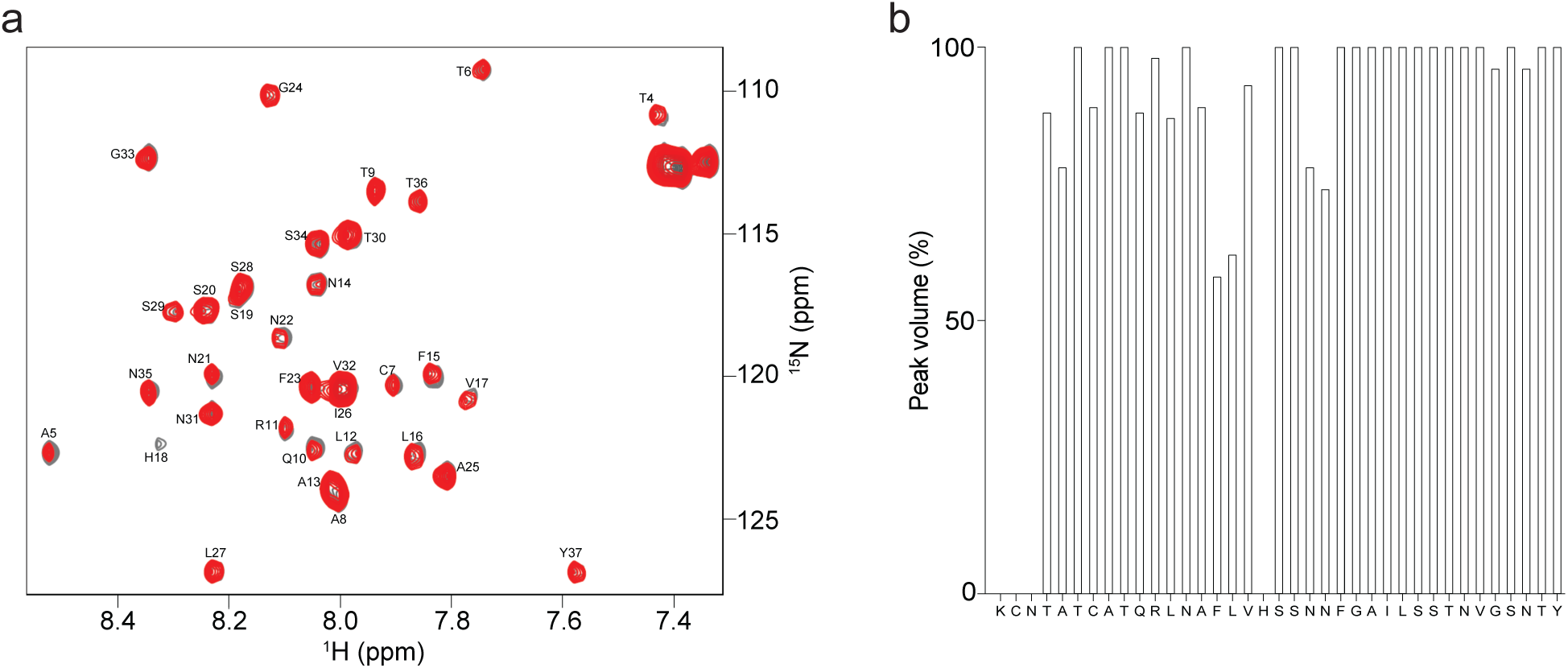
Nuclear magnetic resonance (NMR) spectroscopy characterization of the IAPP-CPP interaction. (a) Superposition of the ^1^H-^15^N heteronuclear single quantum coherence (HSQC) NMR spectra of ^15^N-IAPP before (*grey*) and after (*red*) addition of NCAM1-PrP at an equimolar ratio. (b) Peak volume changes of ^15^N-IAPP residues in the presence of equimolar amount of NCAM1-PrP. As previously reported, the peak volumes for the N-terminal most 3 residues, KCN, could not be determined.^16,20^

Additionally, we performed molecular dynamics (MD) simulations to better understand the observed inhibition of IAPP amyloid self-assembly by the NCAM1-PrP CPP. We focused on dimer formation as it represents a common first step in the amyloid formation pathway.^42,43^ By computing the potential of mean force (PMF) for dimerization – which represents the required free energy change along the reaction coordinate (defined as center of mass distance between the two peptides in the pair) – the interaction strengths of the IAPP–IAPP and IAPP–NCAM1-PrP pairs were compared.

Figure 6 displays the free energy profiles, which revealed a stronger binding interaction of the IAPP–NCAM1-PrP dimer (-15.5 kcal/mol) compared to the IAPP–IAPP dimer (-11.2 kcal/mol). This is likely be due to electrostatic interactions between the negative charge of the aspartic acid residue of the NCAM1 segment of the CPP construct and the three positively charged residues (in particular His18) of IAPP (Table 1). This is in marked contrast to the IAPP–IAPP dimer, where the absence of negatively charged residues means that electrostatic repulsion of the cationic residues is likely to dominate, particularly in the parallel alignment of the peptides that is necessary for amyloid aggregation,^44^ leading to a less energetically favored interaction compared to IAPP–NCAM1-PrP. This result offers a thermodynamics-based explanation for the amyloid inhibition mechanism.

**Figure 6.**
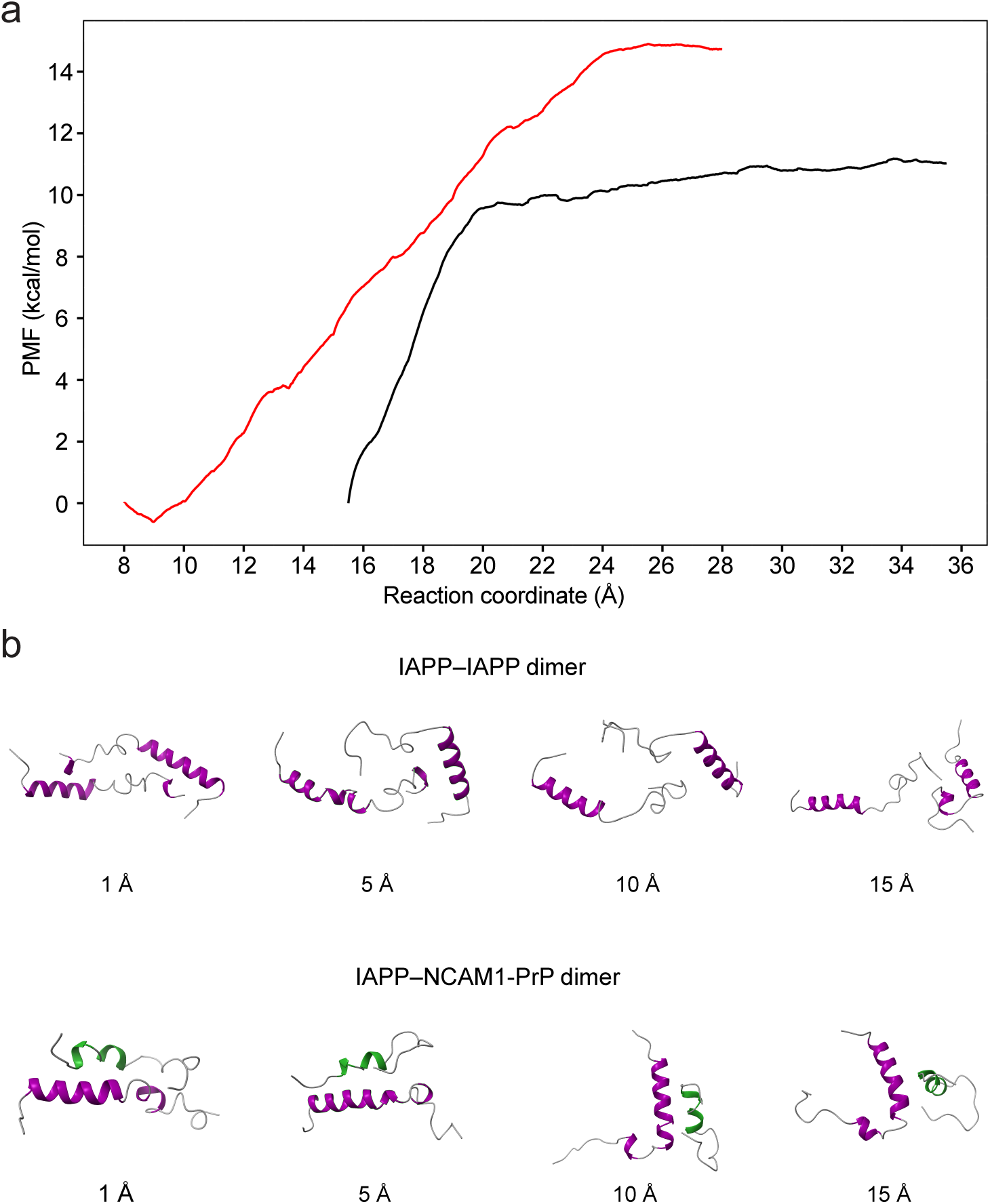
Free energy profiles for the unbinding of the IAPP–IAPP and IAPP–NCAM1-PrP dimeric structures. (**a**) Potential of mean force (PMF) along the reaction coordinate of the IAPP– IAPP (*black*) and IAPP–NCAM1-PrP (*red*) dimers. (**b**) Snapshots of unbinding of the IAPP–IAPP (*top*) and IAPP–NCAM1-PrP (*bottom*) dimers at distances of 1, 5, 10, and 15 Å along the dissociation path. The helical domains of IAPP and NCAM1-PrP are colored purple and green, respectively.

## DISCUSSION

An emerging therapeutic strategy for amyloid diseases is the use of peptides as inhibitors owing to their pharmaceutically-desirable properties, which include greater chemical diversity than other classes of biomolecules, selective binding to specific targets with minimal off-targets, biocompatibility and biodegradability, and ease and low cost of production.^45^ Cell-penetrating peptides (CPPs) are particularly promising as therapeutic agents as they offer an additional advantage in improved delivery to target tissues, cells, and subcellular organelles.^46,47^

In this study, we have used a range of complementary techniques to probe the effects of a designed CPP construct, NCAM1-PrP (Table 1), on IAPP amyloid self-assembly and the associated cytotoxicity. Aggregation assays showed that NCAM1-PrP effectively inhibits IAPP oligomerization and fiber formation (Figure 1). The inhibitory effects of the NCAM1-PrP CPP construct are attributed to the higher stability of the IAPP–CPP interactions compared to IAPP self-interaction (Figure 6 and Table S2). Interestingly, NCAM1-PrP binds to the N-terminal domain, rather than the amyloidogenic central hydrophobic core, IAPP_20–29_, of IAPP (Figure 5 and Table 1). While not amyloid forming in itself, the N-terminal domain of IAPP plays an important role in the peptide’s self-assembly by increasing the nucleation potential of IAPP_20–29_.^10,11^ Of relevance, a range of molecules, including synthetic small molecules and macrocyclic hosts, have been shown to inhibit IAPP amyloid formation by targeting the peptide’s N-terminal domain.^14,16,18–20^ Interestingly, the most prominent interaction of NCAM1-PrP is with a specific residue in the N-terminal domain, namely His18, which has been shown to play a critical role in modulating the membrane interaction, self-assembly and toxicity of IAPP.^48–50^ Thus, NCAM1-PrP binds to the N-terminal domain of IAPP and stabilizes it in a non-amyloid state.

The mechanism of IAPP-induced β-cell cytotoxicity in T2D is yet to be resolved and remains an area of conjecture. Some studies have attributed this cytotoxicity to perturbation of the plasma membrane by extracellular IAPP.^51^ Others have suggested that IAPP toxicity is a consequence of intracellular accumulation of the peptide and disruption of one or more subcellular organelles.^52,53^ Our confocal imaging experiments confirm reports that endocytosis represents one of the cellular uptake routes of IAPP,^50^ which is followed by release of the peptide from endocytic compartments into the cytosol and localization to mitochondria (Figure 3a). Our results are therefore in line with reports that IAPP exerts its cytotoxic effects, at least in part, through disruption of mitochondrial function via direct interaction.^50^ Indeed, mitochondrial damage appears to be a common characteristic of amyloid diseases, with associated proteins – including Aβ, α-synuclein and PrP – directly interacting with, and fragmenting, the organelle.^54–56^

Simultaneous addition of IAPP and NCAM1-PrP to RIN-m cells resulted in strong colocalization of the peptides and inhibition of IAPP cytotoxicity (Figures 2c and 3b). This confirms that the CPPs prevent formation of toxic IAPP oligomers in a cellular environment.

Importantly, time-delayed addition of NCAM1-PrP to IAPP-treated RIN-m cells again resulted in strong colocalization of the peptides of and rescue of IAPP cytotoxicity (Figure 4), demonstrating that the CPP interacts with intracellular IAPP and modulates its toxic structures into non-toxic conformations. Thus, the CPP property of NCAM1-PrP enables it to effectively target both extra- and intracellular IAPP and inhibit formation of its toxic oligomers.

We previously showed that CPPs comprising a hydrophobic signal sequence (SS) fused to an amyloid-derived polycationic NLS-like sequence effectively inhibit conversion of normal cellular PrP (PrP^C^) into the pathogeic scrapie isoform (PrP^Sc^).^21,31^ Subsequently, we demonstrated that these SS-NLS CPPs potently inhibit Aβ oligomerization, fiber formation and neurotoxicity.^22^ Here, we have extended this approach towards T2D by demonstrating that NCAM1-PrP inhibits IAPP amyloid self-assembly and the downstream toxic effects. These studies on diverse amyloid systems strongly suggest that SS-NLS constructs are general amyloid inhibitors, with potential applications in many amyloid-related diseases. The CPP property of these constructs enables them to target both extra- and intracellular amyloid proteins and effectively inhibit their self-assembly and associated cytotoxicity.

## MATERIALS AND METHODS

### Reagents

Purified human IAPP and Cy5-labeled IAPP (Cy5-IAPP) were purchased from Anaspec (Fremont, CA). The inhibitor peptide NCAM1-PrP was synthesized in crude form by Selleck Chemicals (Houston, TX) using standard methods. Acetonitrile, chloroform, dimethyl sulfoxide (DMSO) and deuterated dimethyl sulfoxide (DMSO-d6), Dulbecco’s phosphate buffered saline (PBS), sodium phosphate buffer, thioflavin T (ThT), trifluoroacetic acid (TFA), trizma base and Tween 20 were all purchased from Sigma Aldrich (St. Louis, MO). Alexa Fluor 488 NHS Ester, DAPI and the organelle markers – LysoTracker Green (DND-26), LysoTracker Red (DND-99), MitoTracker Green FM and MitoTracker Red FM – were all from Thermo Fisher (Waltham, Massachusetts, USA).

### Peptide preparation

To remove any pre-formed aggregates, 1–2 mg of IAPP was dissolved in 8 M guanidine hydrochloride (GuHCl), filtered through a 0.2 μm syringe filter, and loaded onto a C-18 microspin column (AmiKa/Harvard Bioscience, Holliston, MA). The column was washed with 10% acetonitrile/0.1% TFA, followed by MilliQ water, and then eluted with 50% acetonitrile/0.1% TFA. The resulting IAPP solution was aliquoted into 0.5 mL Protein LoBind tubes (Eppendorf), lyophilized overnight, and then stored at -80 °C. A fresh stock solution of IAPP was prepared in MilliQ water for each experiment, with the concentration determined by absorbance measurements at 280 nm (ε = 1,280 M^-1^ cm^-1^ for tyrosine). NCAM1-PrP was purified in-house using reverse-phase HPLC. Following purification, the CPP was lyophilized overnight and stored at -80 °C. To prepare the NCAM1-PrP stock solution, the lyophilized peptide was resuspended in MilliQ water and then filtered through a 0.2 μm syringe filter into a 0.5 mL Protein LoBind tube, and stock solution concentration was determined by absorbance measurements at 280 nm (ε = 5,690 M^−1^ cm^−1^ for tryptophan).

### NCAM1-PrP labeling

The NCAM1-PrP CPP construct was N-terminally labeled with Alexa Fluor 488 (Alexa_488_) NHS ester following a published protocol.^22^ Briefly, ∼0.5 mg lyophilized peptide was dissolved in 8 M GuHCl, filtered and loaded onto a C-18 microspin column. The column was then washed (10% acetonitrile/0.1% TFA followed by MilliQ water) to remove unbound peptide, and a dye solution (0.1 mg Alexa_488_ in 0.1 M NaHCO_3_, pH 8.4) was added to the column, which was vortexed and incubated on a rotor for 2–4 h at room temperature. The column was washed again to remove unbound dye, and the Alexa_488_-labeled peptide (A_488_-NCAM1-PrP) was eluted with DMSO. Labeling efficiency was determine by mass spectrometry (QToF LC/MS) and concentration of labeled peptide was measured using absorbance at 494 nm (ε = 73000 M^-1^cm^-1^).

### Thioflavin T (ThT)-based kinetic aggregation assay

The aggregation kinetics of IAPP in the absence or presence of NCAM1-PrP were measured in black 96-well, flat bottom, plates (Corning Inc.; NY) using a Synergy H1MF Multi-Mode microplate reader (BioTek; Winooski, Vermont). For each experiment, three solutions were prepared and kept on ice: A (freshly prepared IAPP in MilliQ water), B (freshly prepared NCAM1-PrP in MilliQ water), C (ThT dissolved in MilliQ water). Then, solutions A and/or B were mixed with C in PBS to produce solutions with the following final concentrations: 10 μM IAPP, 0–10 μM NCAM1-PrP, and 40 μM ThT. Additionally, a solution of 40 μM ThT in PBS was prepared to be used as a blank for normalization. Peptide aggregation was monitored by 10 second linear shaking and measuring the ThT fluorescence (λ_ex/em_ = 440/480 nm) at 10-min intervals at 37°C.

### Transmission electron microscopy (TEM)

Solutions of 10 μM IAPP, alone or with an equimolar amount of NCAM1-PrP, were prepared and incubated for 24 h at 37 °C. Thereafter, a 5 μL droplet of each solution was placed on a freshly glow-discharged copper/formvar/carbon grid (400 mesh, Plano Gmbh). After 5 min, the droplets were wicked away by placing the grid perpendicularly on a filter paper. This was followed by gently washing the grid by dipping the upper surface in a water droplet and drying it using a filter paper as described above. The grid was then placed upside down for 15 s in a 1% uranyl acetate solution (Electron Microscopy Sciences; Hatfield, PA). Finally, the grid was washed thrice with water, dried for 10 min under cover at room temperature, and imaged on a Talos F200X TEM (FEI; Hillsboro, OR) equipped with a Ceta 16M camera operated at an accelerating voltage of 200 kV. Images were processed and analyzed using the Fiji image processing software.^57^

### Dot blot assay

Samples of 10 μM IAPP, alone or with an equimolar amount of NCAM1-PrP, were incubated for 0–6 h. The peptide solutions were then applied to a nitrocellulose membrane and dried for 1 h at room temperature or overnight at 4 °C. The membranes were subsequently blocked with 5% non-fat milk in TBST buffer (20 mM Tris, 0.01% Tween 20; pH 7.4) for 1 h at room temperature, washed thrice with TBST buffer and incubated overnight at 4 °C with polyclonal A11 antibody (1/1000 dilution in 5% non-fat milk in TBST buffer; Life Technologies Corp.; Grand Island, NY). Next, the samples were washed thrice with TBST buffer and incubated with HRP-conjugated anti-rabbit IgG (1/500 dilution in 5% non-fat free milk in TBST buffer; Santa Cruz Biotechnology; Dallas, TX) at room temperature for 3 h. Subsequently, the dot blots were washed thrice with TBST buffer, developed using the ECL reagent kit (Amersham, Piscataway, NJ), and finally imaged on a Typhoon FLA 9000 instrument (GE Healthcare Life Sciences; Pittsburgh, PA) with the settings for chemiluminescence. The chemiluminescence signal intensity was quantified using the Fiji software.

### Cell culture

Rat insulinoma RIN-m cells (ATCC no. CRL-2057; Manassus, VA) were cultured in RPMI-1640 medium supplemented with 10% FBS (GE Healthcare Life Sciences; Logan, UT), 1% penicillin/streptomycin and 0.5 mM 2-mercaptoethanol (both from Sigma), in 5% CO_2_ at 37 °C. Upon reaching 95% confluence, the cells were split with 0.25% trypsin-EDTA (Sigma) into fractions that were propagated or used in experiments.

### Cell viability assay

RIN-m cells were plated at a density of 5 × 10^3^ cells/well in 100 μL complete medium in 96-well plates and cultured in 5% CO_2_ at 37 °C for 24 h before removing the medium and replacing it with 90 μL of FBS-free media. 10 μL of solutions containing IAPP, alone or co-mixed with NCAM1-PrP, at the indicated concentrations, were then added to the wells and incubated for the indicated durations in 5% CO_2_ at 37 °C. For the time-delayed addition experiments, cells were exposed to IAPP for 6–24 h before addition of NCAM1-PrP and incubation of the cells for a total duration of 72 h. Subsequently, 20 μL MTS reagent was added to each well and incubated for 4 h in 5% CO_2_ at 37 °C. Absorbance of the soluble formazan product (λ = 490 nm) of MTS reduction was measured on a Synergy H1MF Multi-Mode microplate reader, with a reference wavelength of 650 nm to subtract the background. Wells treated with peptide-free carrier served as a control. Cell viability was determined from the ratio of the treated wells to the control wells.

### Confocal fluorescence microscopy

RIN-m cells were plated at a density of 3 × 10^5^ cells/well in 300 μL complete medium in μ-slide 8-well (ibidi GmbH; Gräfelfing, Germany) and cultured for 48 h. Thereafter, the medium was replaced with fresh FBS-free medium containing 5μM IAPP (IAPP:Cy5-IAPP, 4:1 molar ratio) or 5μM A_488_-NCAM1-PrP and incubated for 24 h. For co-staining of intracellular organelles, 10 min prior to imaging the medium was replaced with fresh FBS-free medium containing organelle markers (DAPI and 200 nM MitoTracker Red FM/MitoTracker Green FM or 50 nM LysoTracker Red DND-99/LysoTracker Green DND-26) was added. For peptide co-localization experiments, the cells were treated with 5μM IAPP (IAPP:Cy5-IAPP, 4:1 molar ratio) co-mixed with 5μM A_488_-NCAM1-PrP for 24 h. For the time-delayed addition experiments, the cells were treated with 5μM IAPP (IAPP:Cy-5 IAPP, 4:1 molar ratio) for 24 h, which was followed by addition of 5 μM A_488_-NCAM1-PrP and incubation of the cells for a further 48 h. Finally, the medium was removed, the cells were washed with PBS to remove any extracellular peptides or organelle markers, and 300 μL fresh FBS-free medium was added. Images were acquired on a Leica Stellaris 8 confocal microscope equipped with a 63×Plan-Apo/1.3 NA oil immersion objective with DIC capability, and images were processed using the Fiji image processing software.

### Nuclear magnetic resonance (NMR) spectroscopy

Recombinant human IAPP was produced using a cleavable fusion construct as described.^58^ Two-dimensional HSQC NMR experiments were performed on a 600 MHz Bruker instrument at 12 °C. The NMR sample (350 µL) contained ^15^ N-IAPP at a concentration of 25 µM in 20 mM sodium phosphate (pH 6.2), with 10% D_2_O in a Shigemi NMR tube (Shigemi Inc.; Allison Park, PA). A stock solution of 10 mM NCAM-PrP was prepared in DMSO-d_6_. For each NMR experiment, a freshly prepared aliquot of ^15^ N-IAPP was used to avoid potential complications from amyloid formation. NMR spectra were processed using the MNova software.

### Molecular modeling

***System preparation.*** The starting structure of the monomeric form of IAPP was generated from the PDB entry 5MGQ,^59^ which has 27% secondary structure. The initial structure of the monomeric form of NCAM1-PrP was constructed from the peptide’s primary sequence, with a deprotonated Asp residue and protonated Arg and Lys residues, to fulfill the physiological pH condition, while the N-and C-termini were capped with acetyl (ACE) and N-methyl (NME) groups, respectively, to maintain their neutrality, using UCSF ChimeraX (version 1.8).^60^

The LEaP module of Amber24^61^ was used for adding missing atoms, applying the ff19SB protein force field,^62^ solvating the proteins in a truncated octahedron box of OPC water molecules,^63^ with a buffering distance set to 12.0 Å, loading Li/Merz ion parameters (12-6-4 set) for monovalent ions,^64^ in the designated water model, and neutralizing the modeled system with charge neutralizing counter ions (i.e. Cl^−^). For the dimeric forms, K^+^ and Cl^−^ ions (at a concentration of 0.15 M) into the water model to mimic a typical biological environment.

***Simulated annealing.*** Minimization, heating, and simulated annealing were performed on the system using the PMEMD engine within Amber24. For the NCAM1-PrP monomer, three minimization stages were conducted. Each minimization process consisted of steepest descent and conjugate gradient energy minimization processes. The first minimization stage included 10,000 cycles with positional restraint using a force constant of 20.0 kcal/(mol·Å^2^) on all peptide atoms. The second minimization stage included 5,000 cycles with positional restraint using a force constant of 1.0 kcal/(mol·Å^2^) on all peptide backbone atoms. The third minimization stage included 5,000 cycles. The applied positional restraints were relative to the initial coordinates of the modeled system.

After the minimization stages, a heating process was conducted on the system for 0.5 ns, starting from 0 K up to the physiological temperature of 310 K. The heating process used a constant volume condition, a time step of 1 fs, Langevin dynamics^65^ with a collision frequency of 5.0 ps^−1^, non-bonded cutoff of 8.0 Å, SHAKE algorithm^66^ to perform bond length constraints on hydrogen atoms, and the Particle-Mesh Ewald (PME) method^67^ with its default parameters.

To obtain unstructured conformation of the NCAM1-PrP monomer, nine stages of simulated annealing was performed after conducting the minimization and heating processes. In each stage and for 10 ns, the system was quickly heated from 310 K up to 1010 K, then slowly cooled down to 310 K. A random conformation with 32% secondary structure was achieved with this protocol.

***Dimer structures.***To obtain IAPP–IAPP dimer, the monomeric form of IAPP was docked on another copy of itself, whereas the IAPP–NCAM1-PrP dimer was obtained by docking the monomeric form of IAPP on the monomeric form of NCAM1-PrP. The docking studies were carried out using the HADDOCK web server.^68,69^ Bound conformations were predicted and assessed using the HADDOCK score and Z-Score.

***Molecular dynamics simulations.*** Molecular dynamics (MD) simulations were performed on the dimeric systems using the PMEMD engine within Amber24. The total preparation simulation time for each system with positional restraint was 11.2 ns (Table 2). The system preparation protocol included two energy minimization processes (100 cycles were steepest descent energy minimization, and the rest of the cycles were conjugate gradient energy minimization), heating from 0 to 310 K, and several equilibration processes at different periodic boundary conditions. The applied positional restraints were relative to the initial coordinates of the modeled system.

**Table 2.** Summary of the preparation protocol for simulations of the dimeric systems.

| Process | Ensemble<sup>a</sup> | Positional restraint [$k_f$]<sup>b</sup> | | Steps |
| --- | --- | --- | --- | --- |
|  |  | Whole peptide | Peptide backbone |  |
| Minimization 1 | NVT | 20.0 |  | 10,000 |
| Minimization 2 | NVT |  | 20.0 | 10,000 |
| Heating<sup>c</sup> | NV |  | 20.0 | 500,000 |
| Equilibration 1<sup>c</sup> | NVT |  | 20.0 | 100,000 |
| Equilibration 2<sup>c</sup> | NPT |  | 20.0 | 100,000 |
| Equilibration 3<sup>c</sup> | NPT |  | 15.0 | 100,000 |
| Equilibration 4<sup>c</sup> | NPT |  | 10.0 | 100,000 |
| Equilibration 5<sup>c</sup> | NPT |  | 5.0 | 100,000 |
| Equilibration 6<sup>d</sup> | NPT |  | 1.0 | 100,000 |
| Equilibration 7<sup>d</sup> | NPT |  | 0.1 | 2,500,000 |
| Equilibration 8<sup>d</sup> | NPT |  |  | 2,500,000 |
| <sup>a</sup> N: constant number of atoms; V: constant volume; T: constant temperature; and P: constant pressure.<br><sup>b</sup> $k_f$: force constant in units of kcal/(mol·Å<sup>2</sup>).<br><sup>c</sup> Time step was assigned to 1 fs.<br><sup>d</sup> Time step was assigned to 2 fs. | | | | |

The temperature was maintained at physiological value of 310 K using Langevin dynamics with a collision frequency of 5 ps^−1^. The pressure was maintained at 1 bar with isotropic position scaling using Berendsen barostat^70^ with a pressure relaxation time of 1 ps. The non-bonded cutoff was assigned to 8.0 Å. The PME method with its default parameters was used to calculate the full electrostatic energy of the unit cell in a macroscopic lattice of repeating images. The SHAKE algorithm was used in all simulation processes (apart from minimization), to constrain hydrogen atoms. Consequently, the time step was assigned to 2 fs for dynamics integration, except at specific processes mentioned in Table 2, where it was assigned to 1 fs to maintain system stability.

***Potential of mean force for the peptide dimers.*** The potential of mean force (PMF), i.e. the free energy profile, of dimer formation can be obtained by conducting adaptive steered molecular dynamics (ASMD).^71,72^ Steered molecular dynamics (SMD)^71^ uses a pseudo particle that puts a steering force on the system in order to cross a reaction coordinate at a particular velocity. In ASMD, the pre-established reaction coordinate is divided into several stages, within which SMD is conducted and the Jarzynski average is computed throughout the stage. For each stage, a single trajectory is chosen according to the one whose work value is the closest to the Jarzynski average. The coordinates at the end of the stage of the selected trajectory are used to start the next stage of SMD trajectories. By engaging the Jarzynski’s equality,^73^ the non-equilibrium work (*W*) done on the system during the SMD simulation can be related to the free energy change (Δ*G*) according to the following equation: Δ𝐺 = −1/𝛽 ln ⟨𝑒^(−𝛽𝑊) ⟩, where 𝛽 = 1⁄(𝑘_*B*_𝑇), *k*_B_ is the Boltzmann constant and *T* is the absolute temperature. The average 〈⋯ 〉 in the equation is taken over the ensemble of SMD trajectories.

Each dimer form was slowly pulled apart along the reaction coordinate, defined as the center of mass (COM) distance between the two peptides. The heavy backbone atoms of the peptide were used to calculate its COM. The restraining force that was applied along the reaction coordinate was 5.0 kcal/(mol·Å^2^). The reaction coordinate was split into 20 stages, each with a length of 1.0 Å. For each stage, 100 trajectories were performed and each trajectory was run for 1.0 ns.

***Data analysis.*** R language (version 4.0.4), was used for composing the analysis script that produced the PMF figure. Molecular visualization and analyses were performed using UCSF ChimeraX (version 1.8).

## ACKNOWLEDGEMENTS

TEM and confocal fluorescence imaging experiments were carried out using the Core Technology Platforms (CTP) resources at NYU Abu Dhabi. NMR experiments were done using instrumentation in the Department of Chemistry at NYU. Molecular modeling was carried out on the High Performance Computing (HPC) resources at NYU Abu Dhabi. This work was supported by funding from NYU Abu Dhabi (grant no. AD389) to M. Magzoub and from NYU to A.D. Hamilton.

## AUTHOR CONTRIBUTIONS

Conceptualization & Project administration, M. Magzoub; Supervision & Funding acquisition, M. Magzoub and A.D.H.; Investigation, Y.O., P.L., M.H., D.M., M. Mustafa, and S.K.; Formal analysis, Y.O., P.L., M.H., D.M., M. Mustafa, S.K., and M. Magzoub; Writing – original draft, Y.O. and M. Magzoub; Writing – review & editing, all authors.

